# A Novel Approach to Study Gamma-Band Coherence in *Ex Vivo* Hippocampal Slices from a Mouse Model of Bipolar Disorder

**DOI:** 10.1101/2021.01.08.425938

**Authors:** Jean C. Rodríguez Díaz, Paul M. Jenkins, Dominique L. Pritchett, Kevin S. Jones

## Abstract

Oscillations play crucial roles in many cognitive processes such as memory formation and attention. GABAergic interneurons can synchronize neuronal activity leading to gamma oscillations (30-60 Hz). Abnormalities in oscillatory activity in the hippocampus have been implicated in the pathology of some mental health disorders including schizophrenia and bipolar disorder, however the neurobiological mechanism underlying these abnormal oscillations are not yet fully understood. We set out to develop a reliable approach to study gamma oscillations in *ex vivo* hippocampal preparations using perforated microelectrode arrays. Perforated microelectrode arrays allow for the simultaneous measurement of electrical activity at multiple sites while allowing solutions to pass through the brain section. We obtained extracellular electrophysiological recordings from acute sections of mouse hippocampus situated on a 60-channel, perforated microelectrode arrays (pMEAs). Bath application of kainate rapidly induced and maintained oscillatory activity in the CA1 and CA3 regions of the hippocampus. Kainate-induced oscillations were quickly abolished by the GABAA receptor antagonist, bicuculline. Furthermore, we employed this approach on a mouse model of bipolar disorder. Sections prepared from mutant mice exhibited an increase in the coherence of gamma power within CA1 despite a reduction in gamma band power.

## Introduction

The brain is composed of networks that communicate through coordinated network activity. In the brain, coordinated networks can give rise to oscillatory activity that can be measured in distinct bands such as theta (4-8 Hz), alpha (8-12 Hz), beta (14-30 Hz) and gamma (30-100 Hz) (G. Buzsáki, 2006; Jensen et al., 2019). Oscillations in the gamma band are crucial for proper execution of many cognitive behaviors including attention, memory and sensory processing (György Buzsáki & Wang, 2012; Cardin et al., 2009; Fries, 2009; Siegle et al., 2014; Tallon-Baudry et al., 1998). Abnormalities in gamma oscillations have been observed in a variety of diseases including schizophrenia, bipolar disorder and Alzheimer’s disease (Goutagny et al., 2013; Gruetzner et al., 2013; Klein et al., 2016; Liu et al., 2012; Nelson et al., 2018; Spencer, 2011; Spencer et al., 2009).

Neuronal networks can be synchronized to generate gamma oscillations by a subset of GABAergic interneurons called parvalbumin-expressing, fast spiking interneurons (FSIs)(György Buzsáki & Wang, 2012; Cardin et al., 2009; Mann & Mody, 2010; Traub et al., 1997; Wang & Buzsáki, 1996; Whittington et al., 2011). In particular, FSIs are poised to contribute to the synchronization of neurons because they have a high degree of connectivity with both excitatory (Packer & Yuste, 2011) and other inhibitory neurons (Fukuda & Kosaka, 2000). They also exhibit many of the biophysical properties required to generate and sustain the high firing rates (Kawaguchi et al., 1987; Kawaguchi & Kubota, 1997) which are crucial for the generation of gamma oscillations. The importance of FSIs in generating gamma oscillations is further supported by multiple computational models such as the Pyramidal Interneuron Network Gamma (PING) and Interneuron Network Gamma (ING) model((Traub et al., 1997; Wang & Buzsáki, 1996); review György Buzsáki & Wang, 2012; Whittington et al., 2011)).

Gamma oscillations increase the metabolic demands of brain tissue (REF). The high oxygen demands can make it difficult to generate and maintain gamma oscillations in *ex vivo* brain slices (Hájos et al., 2009; Kann et al., 2016). Interface chambers increase the oxygen supply by directly exposing brain tissue to highly-oxygenated, humidified air. Interface chambers have been extensively in the study of gamma oscillations *ex vivo* (Buhl et al., 1998; Fisahn, 2005; Tsintsadze et al., 2015). Importantly, gamma oscillations generated in *ex vivo* preparations retain many similarities to their *in vivo* counterparts (Lu et al., 2011) and have proven useful in mechanistic studies of gamma band oscillations. However, a typical disadvantage of interface chambers is that solution exchange is often limited. Thus, pharmacological manipulation of the brain slices can be slow and highly variable. For example, aqueous delivery of the agonists that provide the chronic excitation needed to generate oscillations in *ex vivo* preparations can require from 30 min to 70 minutes of solution exchange to induce oscillatory activity (Lemercier et al., 2017; C. Lu et al., 2012; Pietersen et al., 2014).

Submerged slice preparations are a useful alternative to interface chambers that permit faster solution exchange (C. Lu et al., 2012). Oxygen delivery is typically insufficient to meet the metabolic demands of continuous gamma oscillations, thus submerged slice preparations were typically used to study transient gamma oscillations induced by brief electrical stimulation (Carmeli et al., 2013) or focal application of kainate (Gloveli et al., 2005; McNally et al., 2011). Rapid oscillations in the beta band (∼15 Hz) were generated in submerged slices with the use a specialized chamber to rapidly deliver oxygenated aCSF above and beneath brain slices from young rats (<PND 20) (Hájos et al., 2009). Further technical innovation demonstrated that sustained gamma band oscillations could be generated in submerged slices (Zemankovics et al., 2013; Juzekaeva et al., 2017), and simultaneous recordings of hippocampal local field potential (LFP) and intracellular recordings slices (Zemankovics et al., 2013). Together, these advances significantly expand the study of gamma oscillations in *ex vivo* hippocampal preparations. However, spatial resolution across the hippocampus remains low.

Here, we set out to develop a novel approach to generate and sustain gamma oscillations in *ex vivo* hippocampal slices with high spatial resolution from adult rodents. Our approach combined the use of perforated microarrays and a fast perfusion rate to rapidly deliver oxygenated aCSF and pharmacological compounds *through* the brain slice. The geometry of the microelectrode array allowed simultaneous recording from hippocampal regions CA1 and CA3 which substantially increased the ability to perform detailed analysis of oscillatory coherence within and between hippocampal subfields. To our knowledge, this is the first report to demonstrate the ability to generate and sustain gamma oscillations in a submerged *ex vivo* brain preparations from adult mice with high spatial precision. We demonstrate the power and utility of our approach by characterizing gamma oscillations generated in hippocampal slices from a mouse model of bipolar disorder (Nelson et al., 2018).

## Methods

Mice were housed in the temperature and light-controlled (12/12 h light/dark cycle) animal care facilities at the University of Michigan and given food and water *ad libitum*. All animal procedures were approved by the University of Michigan’s Institutional Animal Care and Use Committees (IACUC) and conformed to NIH Guidelines for animal use. All mice were aged, PND 21-40. C57BL/6J mice were used to develop the assay whereas the knock in *Ank3* W1989R model was used as the mouse model of bipolar disorder (Nelson et al., 2018). A subset of the data in this manuscript appeared in Nelson et al 2018.

### Electrophysiological recordings

Mice were anesthetized with isofluorane and intracardially perfused with ice cold (4°C) modified N-Methy-D-glucamine (NMDG) HEPES artificial cerebrospinal fluid (aCSF) consisting of: 93 mM NMDG 2.5 mM KCl, 0.5 mM CaCl_2_, 10 mM MgCl_2_, 1.2 mM NaH_2_PO_4_, 20 mM HEPES, 25 mM dextrose, 5 mM Ascorbic Acid, 2 mM Thiourea and 3 mM Na-pyruvate. pH was maintained at 7.4 by saturation with O_2_/CO_2_, (95/5%, respectively). Mice were quickly decapitated, and the brain was quickly dissected and transferred to a holding chamber with ice cold NMDG HEPES aCSF. Horizontal hippocampal sections (300 µm thick) were prepared with a Leica VT1200. Sections were bi-laterally hemisected and transferred to an intermediate holding chamber maintained at 33° C for 10-12 min with NMDG HEPES aCSF and then transferred to camber with an aCSF solution consisting of: 126 mM NaCl, 3 mM KCl, 2 mM CaCl_2_, 1 mM MgCl_2_, 1.25 mM NaH_2_PO_4_, 25 mM NaHCO_3_ and 10 mM dextrose at 33° C for 35 min. The pH of the NMDG HEPES aCSF and of aCSF was maintained at 7.4 by saturating the solution with 95%O_2_/5% CO_2_. Sections were then transferred to room temperature for at least 15 min. For the recording session, sections were mounted on perforated multi-electrode array (pMEAs) (Multichannel Systems, Reutlingen, Germany). Sections were secured to the surface of the pMEA surface by using a peristaltic perfusion system (PPS2, Multichannel Systems, Reutlingen, Germany) to create a slight vacuum through the perforations. The secured sections remained submerged in aCSF (29 −31° C, 95% O_2_/5CO^2^%) and were superfused at a rate of 5-7 ml/min. Local field potentials were recorded at 20 kHz using the MEA2100-System (Multichannel Systems, Reutlingen, Germany). Baseline recordings were obtained for 1 hour. Chemically-induced oscillations were evoked by bath application of kainate (400 nM) for 1 hour. Oscillations were abolished by bath application of the GABA_A_ receptor antagonist, bicuculline (10 µM).

### >Gamma oscillation analysis

To determine the power spectrum, we low pass filtered the data at 100 Hz using a IIR Butterworth filter and down sampled to 1 kHz for analysis. Power analysis was done by multi-taper spectrum analysis using a mixture of custom-written MATLAB scripts or using multi-tapered fourier estimation using 5 tapers with the Chronux, an open source Matlab toolbox (*Chronux Home*, n.d.; Mitra & Bokil, 2007). The absolute power in the gamma band was determined by computing the the area under the curve in the frequency band between XX and XX Hertz. The fractional difference in oscillatory power was calculated by (Power_kainate_ -Power_aCSF_)/Power_aCSF_ for during kainate application and (Power_bicuculline+kainate_ -Power_kainate_)/Power _kainate_ for experiments during bicuculline application. Spectrograms were made by convolving the signals with a morelet wavelet as follows: 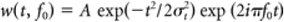, where A is a normalization factor equal to:(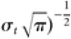 . The width of the wavelet was set to be 25, *m* = *f*_*0*_/σ_*f*_, with σ_*f*_ = 1/2πσ_*t*_. Interhemispheric, zero-phase-lag coherence of gamma oscillations was computed between the LFP on all electrodes on the MEA using the multi-tapered approach above. Coherence was calculated from the autocorrelation on each channel, Sx(f) and Sy(f) and the cross-spectrum between the channels Sxy(f), which is calculated as follows: Cxy(f) = Sxy(f)/sqrt(Sx(f) Sy(f)) (Gregoriou et al., 2009).

Fall time (T_90-10_) and the time to abolish were calculated using MATLAB’s fall time function from the final two minutes of drug application. Fall time was defined as the time required for signal to fall from 90% to 10% of maximum. Time to abolish was defined as the time required for the signal to reach 10% of maximum from the end of kainate application. Recordings obtained from electrodes with artifacts or excessive noise in gamma band were excluded from kinetic analysis. Rise times were calculated by determining the time required to increase the power in gamma band to 90% of maximum, after bath application of kainate or bicuculline, respectively. The power from 25-59 Hz in the final five minutes of the ACSF, the ACSF + 400 nM Kainate and the ACSF + 400 nM Kainate + 10 µM Bicuculline. We calculated peak frequency and the quality factor (Q factor) defined as peak frequency/half bandwidth.

Data were exported to GraphPad Prism for statistical analysis and plotting. Paired t-tests between the kainate and bicuculline application and unpaired t tests between wildtype and mutant mice were done for each brain region in GraphPad Prism. Custom MATLAB scripts are available upon request or at https://github.com/Jcrd25/NeuroMEACode.

## Results

In this study we utilized perforated microelectrode array (pMEAs) chips and perfusion system comprised of two independent perfusion systems to generate and maintain high frequency oscillations in hippocampal sections from adult mice (**Fig 1B**). pMEAs contain a thin, flexible membrane that creates a separate perfusion chamber beneath the microelectrode array and permits aCSF to be delivered above and beneath the slice simultaneously. The first perfusion system pumps warm aCSF directly into the top chamber. A second closed, perfusion system use both a pump and a vacuum to deliver aCSF to the chamber beneath the slice. The pump and vacuum are independently controlled which permits aCSF to be suctioned from the bottom chamber more rapidly than it is delivered. This creates a negative pressure in the bottom chamber that draws aCSF from the top chamber through the brain slice and secures the slice to the array (**Fig 1D**). We oriented each hippocampal section on the rectangular array geometry such that several electrodes were positioned below the CA1 and CA3 hippocampal subfields. Critically, this allowed us to record oscillatory activity from several adjacent locations within each subfield (**Fig 1C**).

**Figure 1:**
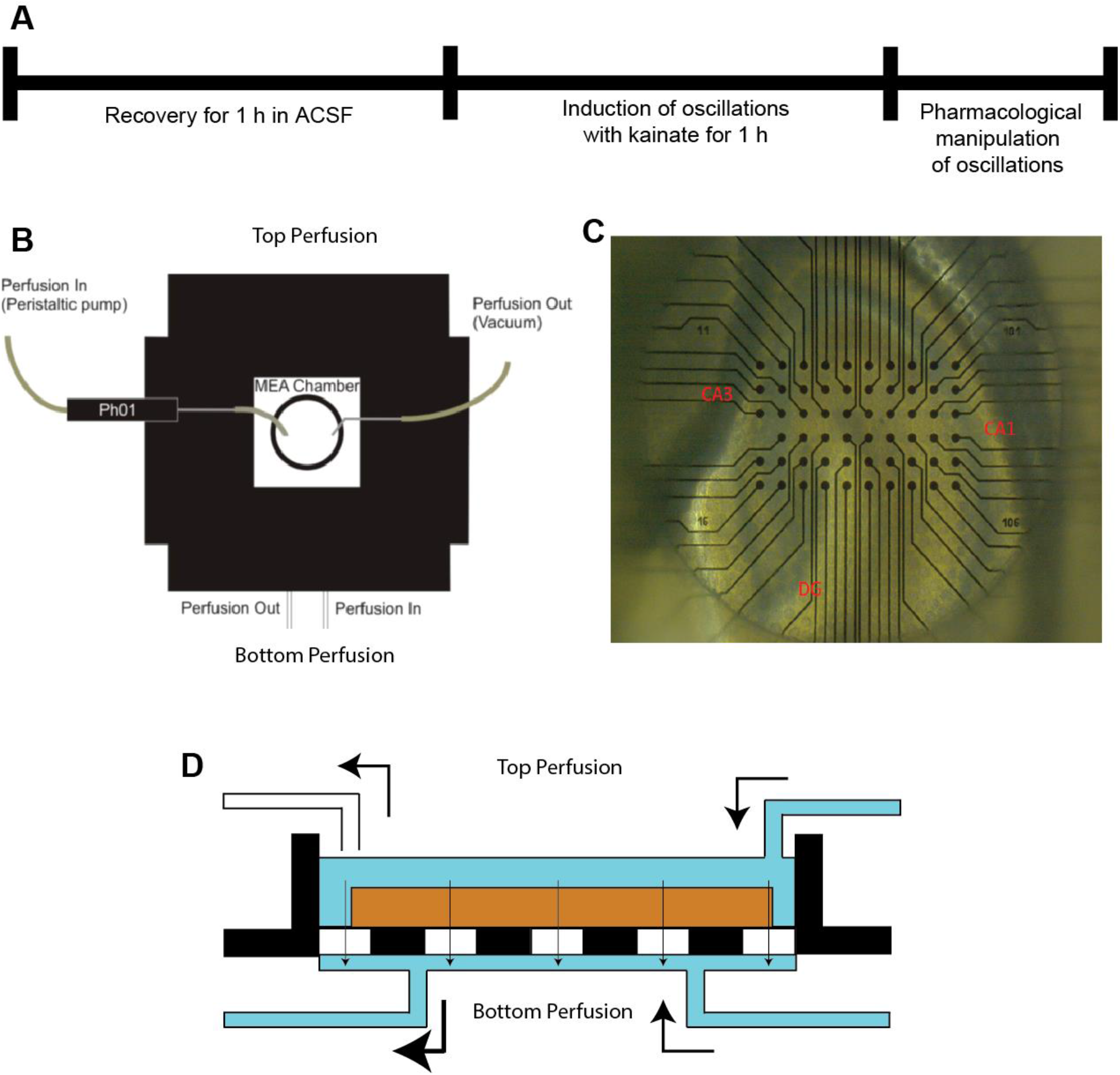
Schematic of Experimental Design and Apparatus. **(A)** Experimental timeline consisted of one-hour acclimation period followed by the perfusion of Kainate 400 mM to induce oscillations and pharmacological manipulations. **(B)** MEA perfusion apparatus. Perfusate is delivered to the MEA chamber via two independent routes. Warm perfusate is delivered to the top chamber of the MEA through an inline heating element (Ph01). Perfusate is delivered to the bottom chamber of the MEA in an independently-controlled, semi-closed loop. Image modified from Multichannel Systems. **(C)** Prototypical layout of the pMEA with a hippocampal slice. Electrode array was arranged to maximize electrode placement in CA1 and CA3 subfields. **(D)** Schematic of the perforated MEA chip. Independently-controlled perfusion systems deliver aCSF above and below MEA chamber. Perfusate is removed from the bottom chamber at a faster rate than it is delivered. The difference in rate creates a small vacuum in the bottom chamber which draws perfusate from the top chamber through the tissue, the perforated membrane and into the bottom chamber. This results in increased diffusion of oxygenated perfusate throughout the depth of the tissue and secures the slice to the MEA.

Bath application of kainate generated visible oscillations in the LFP of both CA1 and CA3. Electrodes in CA1 and CA3 showed little baseline activity (**Fig 2A**). Bath application of kainate induced oscillatory activity in the LFP signal of both CA1 and CA3 (**Fig 2A**. In addition, some slices exhibited a large increase in activity soon after kainate application causing an increase in the power across multiple frequencies which stabilized into oscillatory activity (example can be seen in **Fig6 B,E)**. Kainate application increased the power in oscillations between 20-60 Hz detected in both CA1 and CA3 subfields, suggesting an increase in gamma-band oscillations (**Fig. 2B, C**).

**Figure 2:**
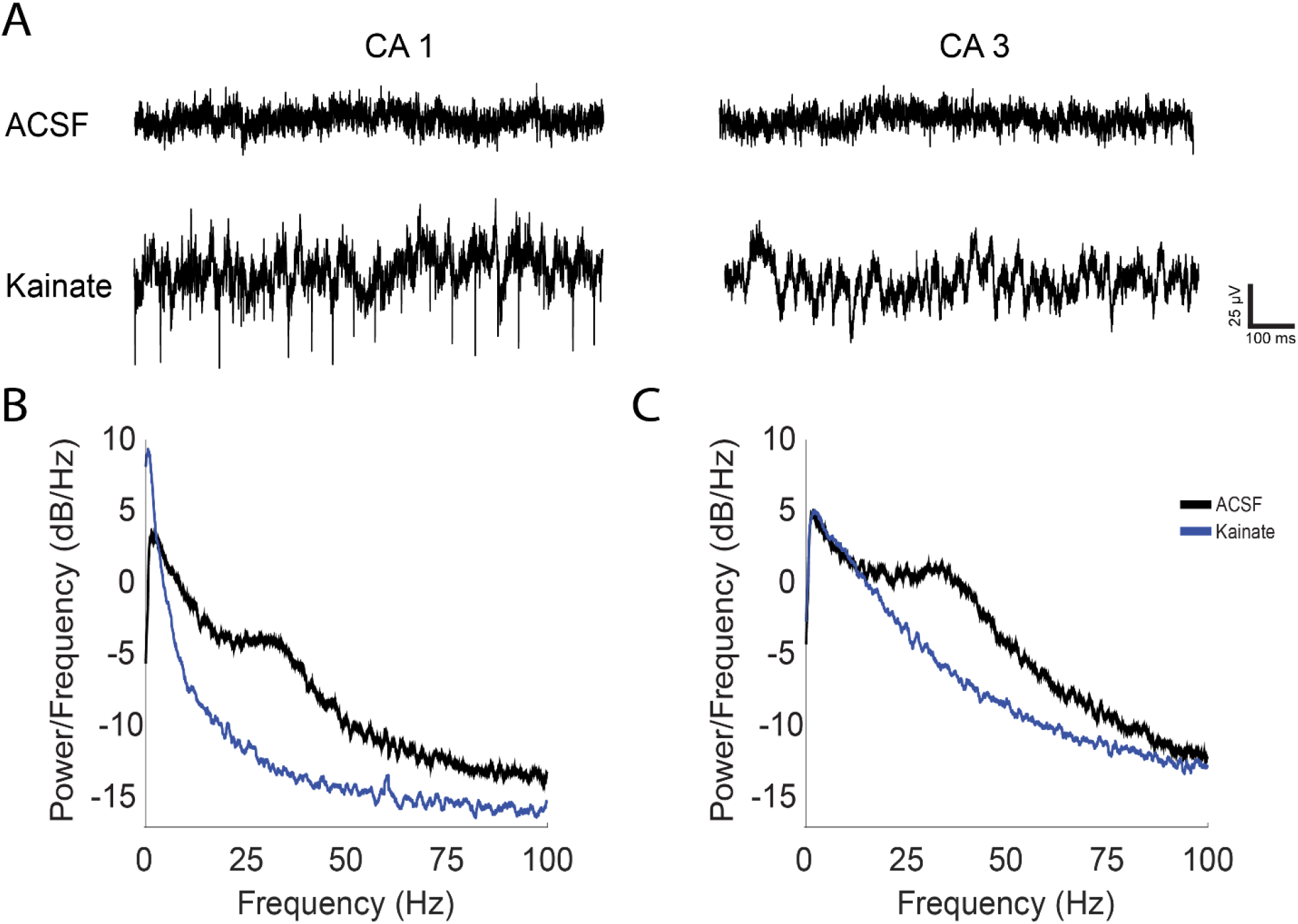
Kainate induces oscillations in CA1 and CA3. **(A)** Prototypical extracellular recordings during ACSF and ACSF with 400 nM kainate application in CA1 and CA3 (low-pass filtered, 100 Hz). **(B and C)** Prototypical periodograms generated from recordings acquired during bath application of vehicle (blue) or kainate (brown) in **(B)** CA1 and **(C)** CA3.

Kainate application induced oscillations in the gamma band in both CA1 and CA3 regions of the hippocampus. Visual inspection of spectrograms revealed that the increase in power in CA1 and CA3 occurred quickly after kainate application and was sustained for the duration of the recording (**Fig. 3A, C)** The peak frequency for the oscillations for both CA1 and CA3 was on the lower end of the gamma band spectrum with the mean CA3 peak frequency being lower than the peak frequency in CA1, even falling into the beta band (**Fig3E**; CA1 30.06 ± 1.149 Hz, N=10, n=93; CA3 21.13 ± 0.9759 Hz, N=10, n=155) unpaired t test p value <0.0001). The quality factor (Q factor) has been used to determine if an LFP signal has periodicity and is oscillating (Lemercier et al., 2017). A Q factor of greater than 0.5 indicates the LFP signal is oscillating. The Q factor of the oscillations in CA1 and CA3 were1.01 ± 0.06,(*N* =10, *n* =93) and 0.74 ± 0.05 (*N* = 10, *n* =155) respectively, indicating that kainate induced oscillations in both CA1 and CA3 (**Fig3F**).

**Figure 3:**
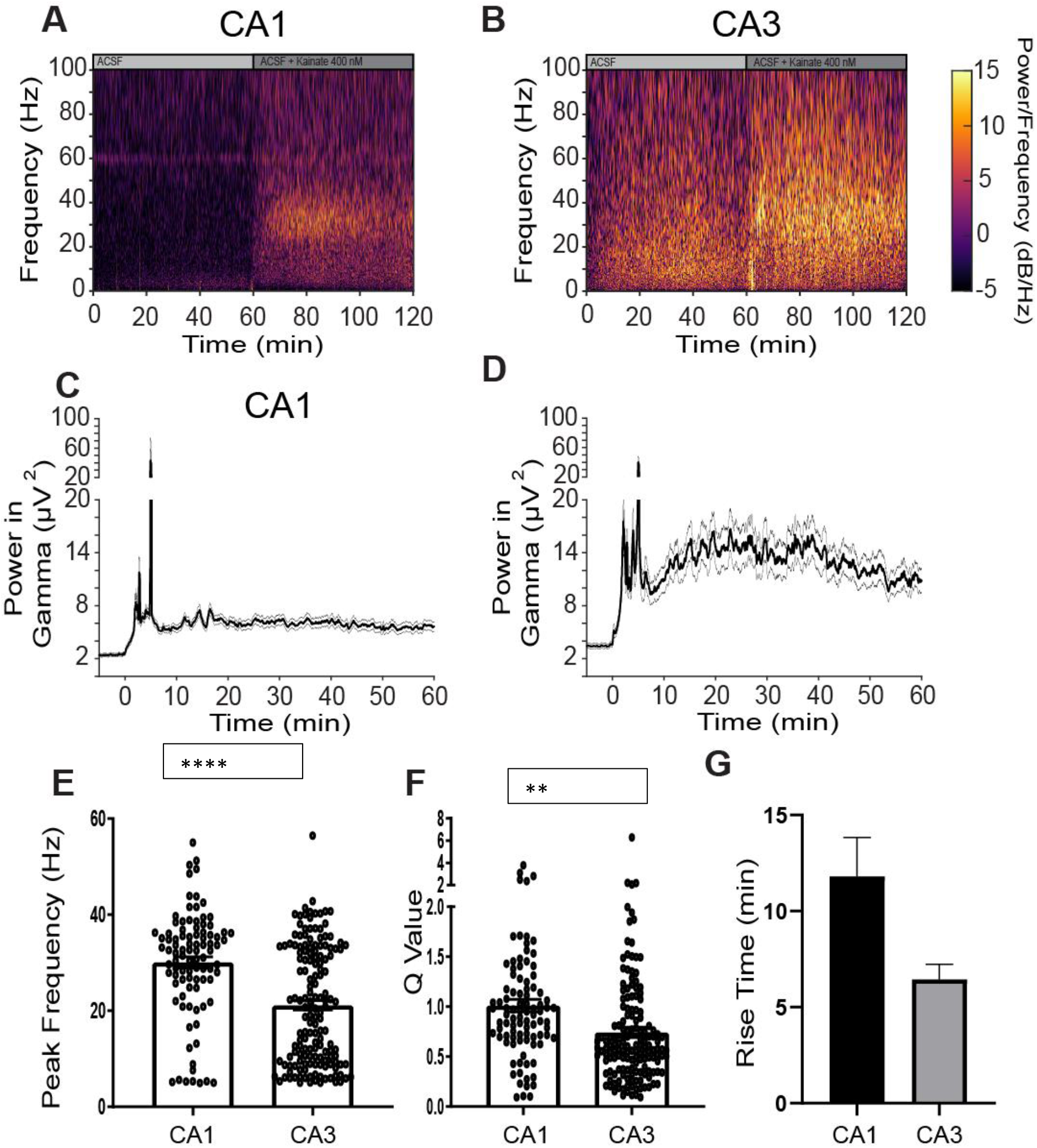
Kainate induces near gamma-band oscillations in CA1 and CA3. **(A, C)** Prototypical spectrograms of a recording from a single electrode in CA1 **(A)** or CA3 **(C)** during bath application of vehicle or kainate (400 nM). **(B, D)** Power in the gamma band (25-59 Hz) from **(B)** CA1 (*N* =10, *n*=93) and **(D)** CA3 (*N*=10, *n*=155). **(E-G)** Quantitative characterization of kainate-induced, gamma-band oscillations. **(E)** Peak frequency (CA1 30.1 ± 1.15 Hz, *N*=10, *n*=93; CA3 21.1 ± 1.0 Hz, N=10, n=155); **(F)** Q value (CA1 1.01 ± 0.06, *N* =10, *n* =93; CA3 0.74 ± 0.05, *N* = 10, *n* =155) and **(G)** Rise time (CA1 ± N=, n=; CA3 ± N=, n=). **** = p value <0.0001;** = p value 0.0013. Unpaired t tests were perfromed in GraphPad Prism.

To determine the role of GABA_A_ neurotransmission, the GABA_A_ receptor antagonist, bicuculline, was co-applied to the bath (**Fig 4**). Bicuculline decreased the gamma-band power of kainate-induced oscillations in both CA1 (fractional difference −0.55 ± 0.03, N = 7, n =71) and CA3 ((−0.59 ± 0.02, N = 7, n =88) (**Fig 4 B, G**). Bicuculline had a quick course and abolished the gamma band power generated by kainate application with a fall time less than two minutes in both CA1 and CA3 (CA1: 62.91 ± 3.1 s, N = 7, n =69; CA3: 69.04 ± 3.6 s, N = 7, n =71; **Fig 4 D, I**). This suggests kainate-induced gamma-band oscillations require GABA_A_ receptors. Moreover, the rapid time course demonstrates the speed of drug delivery in our experimental set-up. From these data, we estimate that fast acting drugs can modulate kainate-induced gamma oscillations in about 70 s after they solutions are switched (**Fig4E&J**).

**Figure 4:**
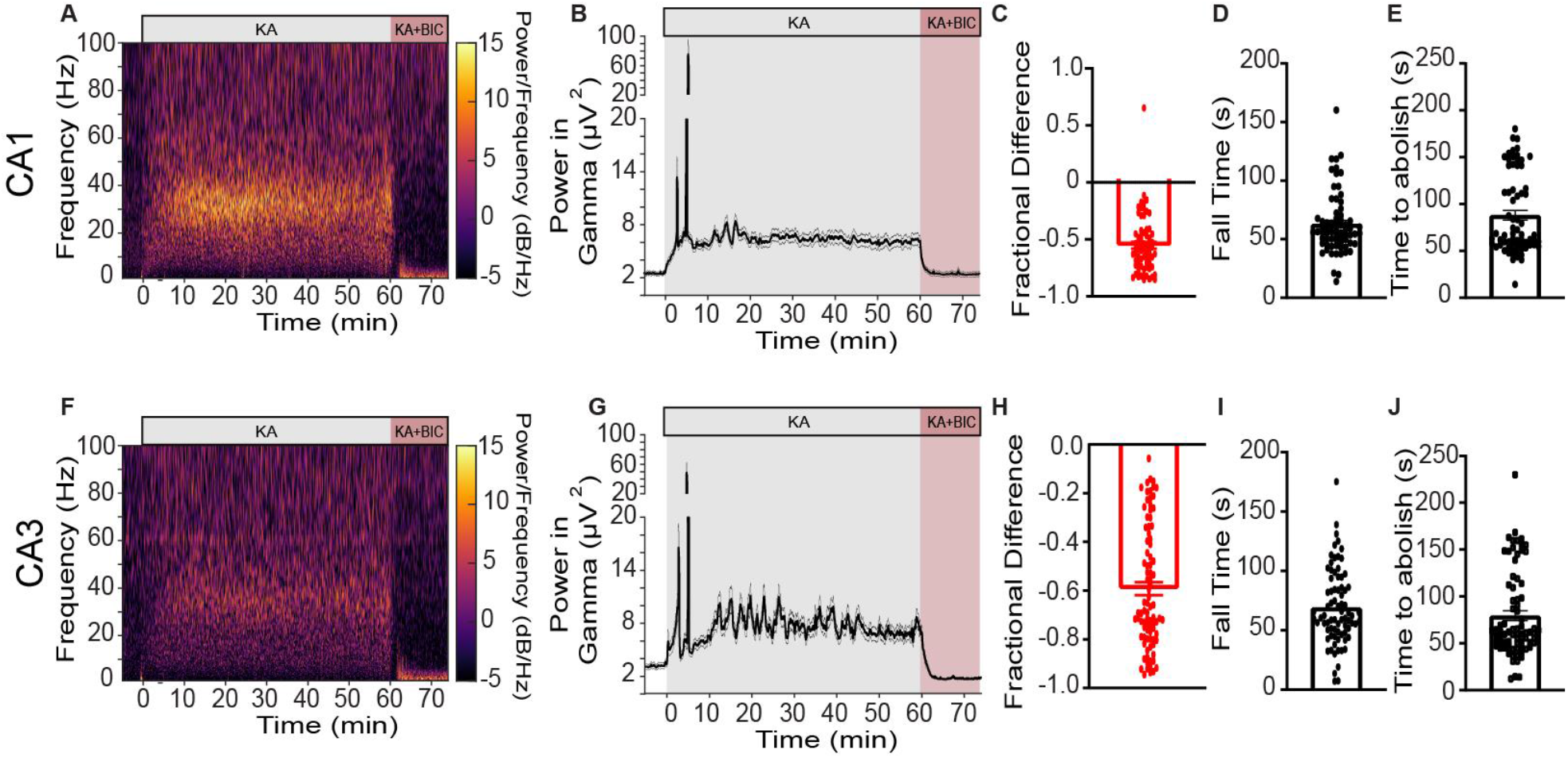
Kainate-induced oscillations are GABA_A_-dependent. **(A, F)** Prototypical spectrogram of individual electrode from **(A)** CA1 and (**F)** CA3. **(B, G)** Power in gamma-band oscillations during wash-in of kainate (400 nM) or kainate and bicuculline (10 µM) (**B)** CA1 (*N* = 7, *n* =71) and **(G)** CA3 (*N*=7, *n*=88). **(C-E) Characterization of Bicuculline phase in CA1. (C)** Fractional difference in power after bicuculline application (−0.55 ± 0.03, N = 7, n =71), **(D)** Fall time (62.91 ± 3.1 s, N = 7, n =69) and **(E)** Time to abolish (88.1 ± 5.1 s, N = 7, n =69). **(H-J) Characterization of Bicuculline phase in CA3. (H)** Fractional difference in power from last 5 minutes of kainate phase to the last 5 minutes of kainate and bicuculline phase (−0.59 ± 0.02, N = 7, n =88), **(I)** Fall time (69.04 ± 3.6 s, N = 7, n =71) and **(J)** Time to abolish (79.6 ± 5.2 s, N = 7, n =77).

Planar MEA chips allow for the simultaneous recording of adjacent regions. A primary advantage of our approach is that the geometry of the electrodes on our MEA chips allowed us to record from several regions within the CA1 and CA3 subfields. This allowed us to determine how the coherence in gamma power changed within *and* between CA1 and CA3 during kainate induced oscillations. Kainate increased the coherence in the gamma band (**Fig5A-E**) of electrodes located within the same subfield more robustly than electrodes located in different subfields regions **(Fig 5f**).

**Figure 5:**
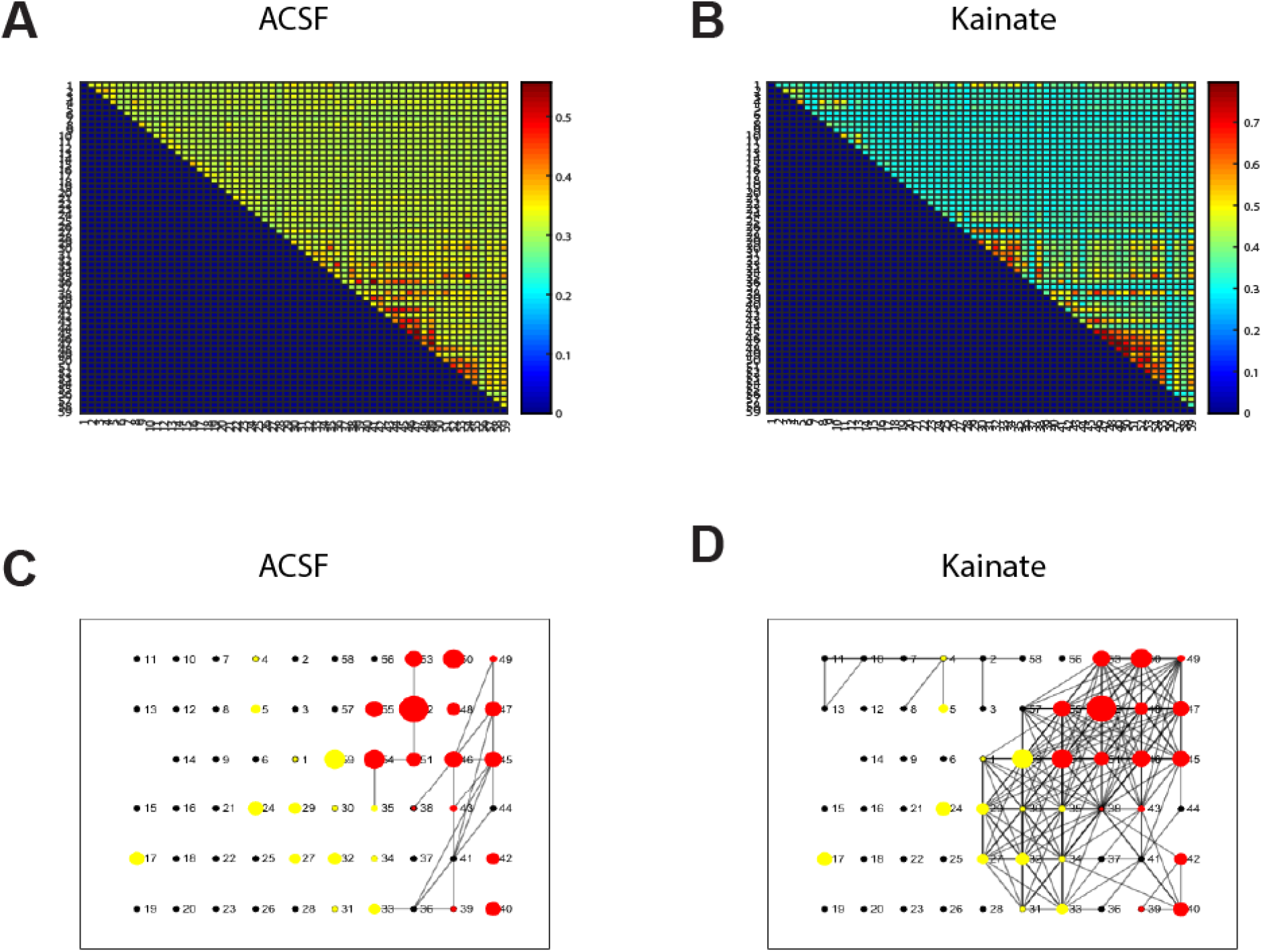
Kainate increases coherence across CA1 and CA3. **(A, B)** Representative correlation map of power of gamma-band oscillation during bath application of vehicle **(A)** or kainate **(B). (C, D)** Representative spatial correlation map during bath application of vehicle **(C)** or kainate **(D)**. CA3 circles are in red, CA1 is yellow. Symbol size is proportional to the power in gamma-band, line represent a correlation strength between two electrodes.

To demonstrate the utility our novel approach we characterize gamma-band oscillations in brain tissue for a mouse model Bipolar Disorder. We utilized the *Ank3* W1989R mice previously reported(Nelson et al., 2018). This mouse line exhibits reduced GABAergic neurotransmission, specifically that mediated by parvalbumin positive FSIs. These mice can generate oscillations in the gamma band in CA1 (**Fig6 A, B**) and CA3 (**Fig D, E**). The power in the gamma band oscillations induced in slices from mutant mice was lower when compared to WT exhibit reduced gamma power in both CA1 and CA3 (**Fig 6C, F**). There was no difference in the fall time between the wildtype and mutant mice in either CA1 (WT: 46.92 ± 4.13 s n= 47; N=5, MT 55.35 ± 6.61 s; n= 59, N =5; p = 0.3108) nor CA3 (WT: 40.48 ± 3.45 s n= 47; N=5; MT 47.61 ± 5.45 s n= 57, N =5; p = 0.2948).

**Figure 6:**
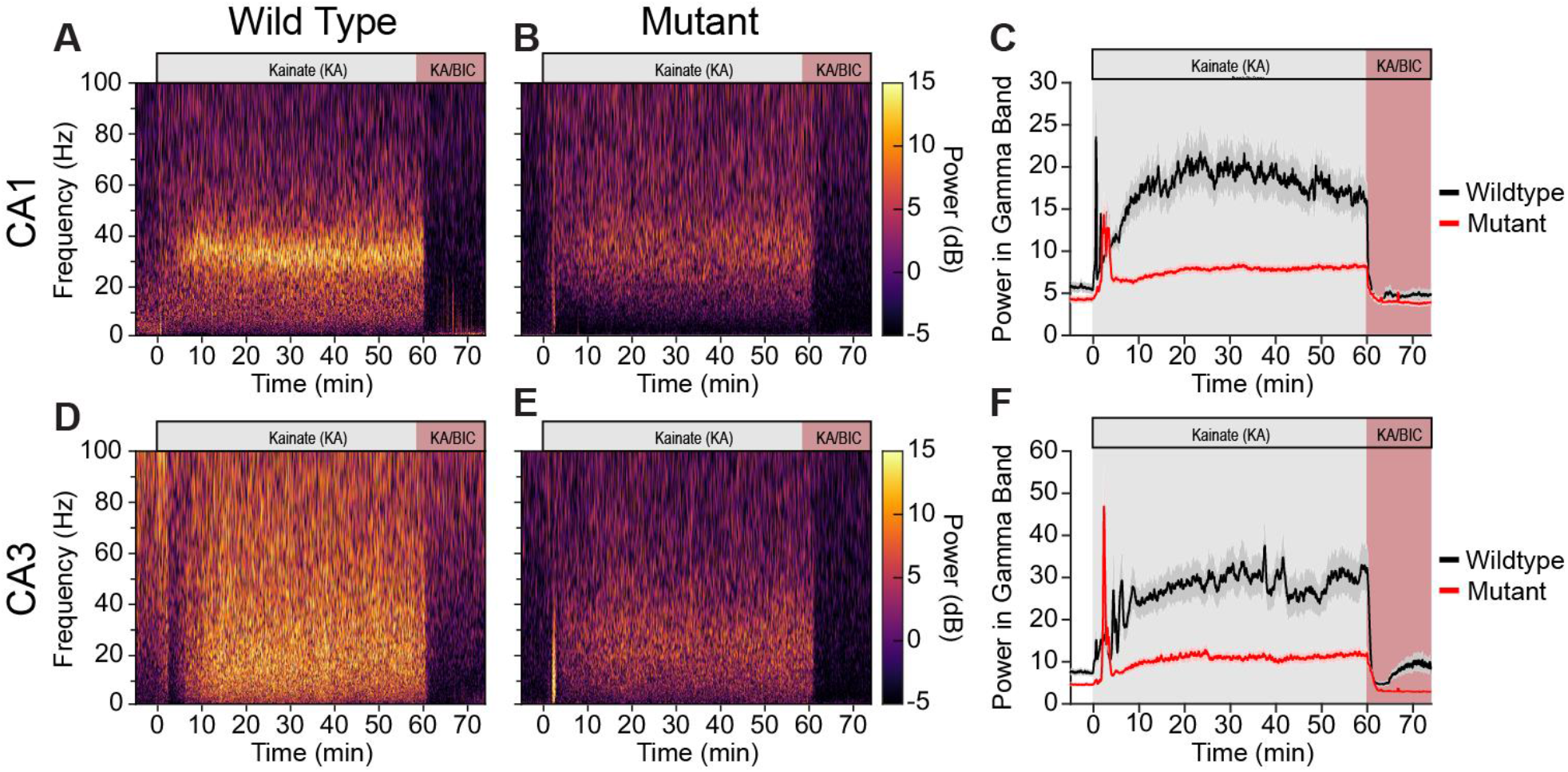
Gamma-band power is reduced in Ank3 mutant mouse. **(A-B)** Representative Power spectrums in CA1 of **(A)** wild type mice and **(B)** mutant mice. **(C)** Power in the gamma band for wildtype mice (black; *N* = 4, *n* =50) and mutant mice (red; *N*=5, *n*=62). Data shown as mean ± SEM. **(A-B)** Representative power spectrums in CA3 of **(D)** wild type mice and **(E)** mutant mice. **(F)** Power in the gamma band for wildtype mice (black; *N* = 5, *n* =50) and mutant mice (red; *N*=5, *n*=59). *N* = number of slices; n = number of electrodes.

Mutant animals exhibited an overall increase in coherence when compared to their wildtype counter parts (**Fig7**). The spatial coherence map of both wildtype and mutant mice show considerable coherence across many of the electrodes, with the mutant mice exhibiting more overall coherence in gamma power (**Fig 7A, B**). Most slices exhibited similar coherence during the aCSF phase within CA1 (WT: 0.410 ± 0.011, N=5; MT 0.402 ± 0.054, N = 4) within CA3 (WT aCSF:0.412 ± 0.010, N=5; MT 0.360 ± 0.017, N = 5) (**Fig7C, D**) with only mutant mice exhibiting an increase in coherence within CA1 (WT: 0.413 ± 0.012, N=5; MT 0.541 ± 0.084, N = 4). Both wildtype and mutant mice exhibited an overall increase in coherence in gamma power within CA3 (WT aCSF:0.412 ± 0.010, N=5; MT 0.360 ± 0.017,N = 5; aCSF+Kainte: 0.506 ± 0.034, N=5; MT 0.528 ± 0.039, N = 4; **Fig7D**), while the coherence across CA1 and CA3 seemed to have slightly decreased in wildtype animals but not change in the mutant mice (WT aCSF: 0.395 ± 0.010, N=5; MT 0.327 ± 0.007,N = 4; aCSF+Kainate: 0.359 ± 0.009, N=5; MT 0.340 ± 0.014,N = 4).The change in coherence after kainate wash in was higher within CA1 (WT: 1.008 ± 0.025, n =5, MT: 1.175 ± 0.093, n=4; p = 0.0065) and across CA1 and CA3 (WT: 0.908 ± 0.018, n=5, MT: 1.044 ± 0.057, n=4, p = 0.0417) in the mutant animals when compared to their wild type counterparts (**Fig 7F**). They also exhibited stronger coherence across CA1 and CA3 regions (**Fig7F**).

**Figure 7:**
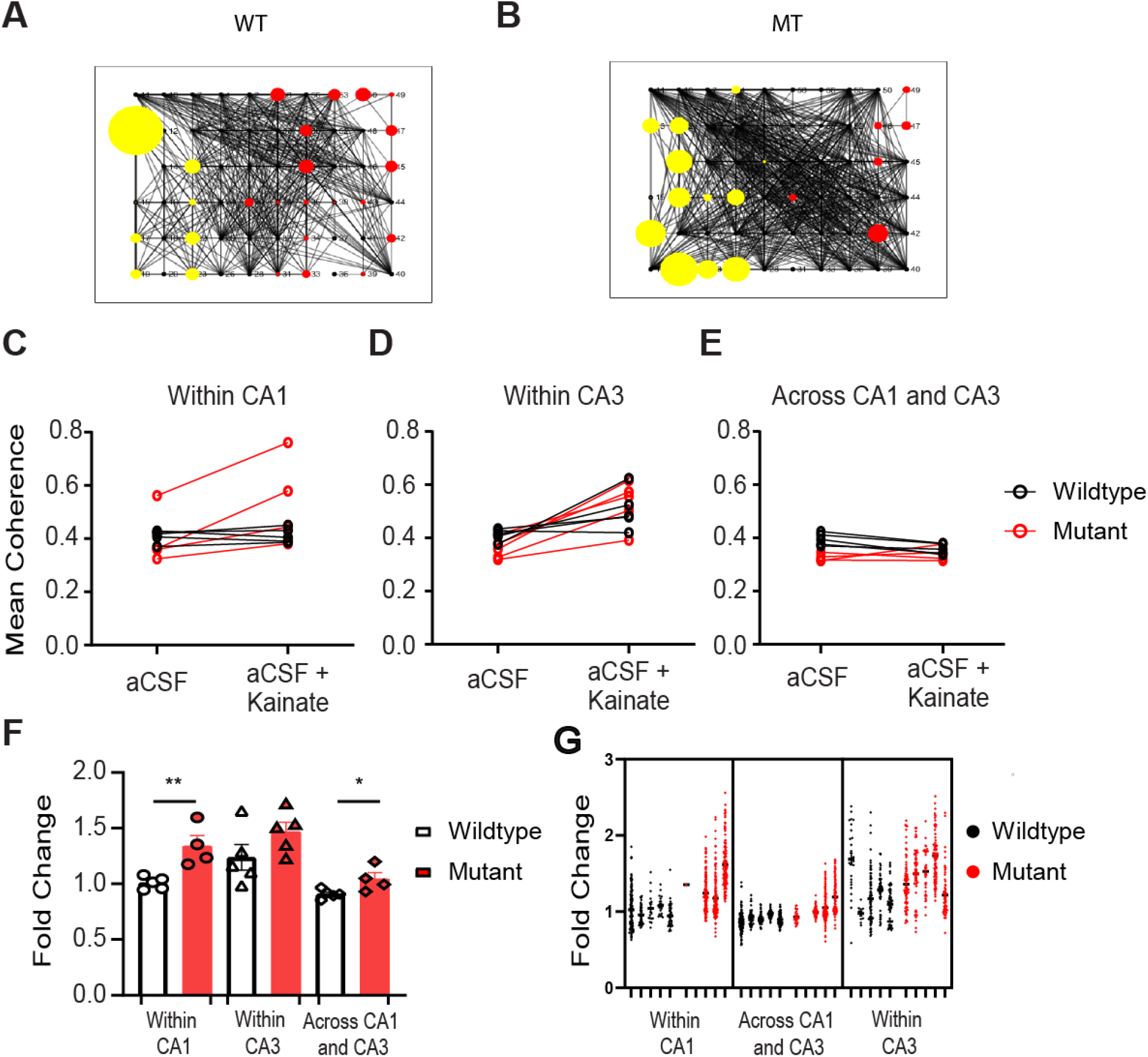
Coherence increases in the mutant mouse model. **(A, B)** Representative map of spatial correlation during bath application of kainate for **(A)** wiltype and **(B)** mutant mouse. Red circles indicate electrodes positioned in CA1; yellow circles indicate electrodes positioned in CA3. Circle diameter represents proportion of gamma-band power; line width indicates strength of correlation between electrodes. **(C-E)** Mean coherence during the last five minutes of aCSF and during aCSF + Kainate for electrode pairs **(C)** within CA1 (WT aCSF:0.410 ± 0.011, N=5; MT 0.402 ± 0.054,N = 4; aCSF+Kainte: 0.413 ± 0.012, N=5; MT 0.541 ± 0.084,N = 4), **(D)** within CA3 (WT aCSF:0.412 ± 0.010, N=5; MT 0.360 ± 0.017,N = 5; aCSF+Kainate: 0.506 ± 0.034, N=5; MT 0.528 ± 0.039, N = 4), and **(E)** across CA1 and CA3 (WT aCSF: 0.395 ± 0.010, N=5; MT 0.327 ± 0.007,N = 4; aCSF+Kainate: 0.359 ± 0.009, N=5; MT 0.340 ± 0.014,N = 4). **(F)** Fold change in mean coherence from aCSF to aCSF +Kainate. **(F)** Bar graph of the mean coherence within CA1(WT: 1.008 ± 0.025, n =5, MT: 1.175 ± 0.093, n=4; p = 0.0065), within CA3 (WT: 1.238 ± 0.115, n=5, MT: 1.468 ± 0.084, n=5; p = 0.1451) and across CA1 and CA3 (WT: 0.908 ± 0.018, n=5, MT: 1.044 ± 0.057, n=4, p = 0.0417). Data shown as Mean ± SEM. * = p value 0.0417;** = p value 0.0065, n = number of slices. Unpaired t tests were perfromed in GraphPad Prism. **(G)** Fold change in coherence for each electrode comparison grouped in slices. Individual circles represent a correlation in the gamma band between two electrodes. Black circles represent wildtype and red circles represent mutant mice.

## Discussion

In this study, we demonstrated a novel approach for studying the coherence of kainate-induced gamma oscillations in acute hippocampal slices.

We found that coherence within regions was stronger than across regions. We successfully employed this approach on a mouse model of bipolar disorder that allowed us to quantify that gamma oscillations were lower in power in the mutant animals, but they exhibited a larger fold changes in coherence within CA1 and across CA1 and CA3 when kainate induced gamma oscillations, but not within CA3. We also found that both gamma oscillations from mutant and wildtype mice were GABA A receptor dependent with similar fall times. Taken together this suggest that the resulting deficits in FSI neurotransmission caused by the Ank3 mutation may lead to an overall decrease in the power in the gamma band while potentiating the coherence in the gamma power.

These findings are largely a result of combining several technical innovations, particularly the use of pMEAs to generate rapidly generate, sustain, and pharmacologically manipulate gamma oscillations in *ex vivo* hippocampal sections. Overall, our results are supported by previous reports in which bath application of excitatory chemicals induce gamma oscillations in *ex vivo* hippocampal slices. However, the approach we use in our study is distinct in that it expands previous studies of gamma oscillations in two novel and important parameters. Firstly, the density and the geometry of the electrode arrays used in this study were chosen to maximize the number of simultaneous recordings that could be acquired within each subfield. Previous reports generally place one electrode in CA3, whereas each hippocampal slice used in this study had an average of 10 electrodes placed in CA1 and 13 placed in CA3. The increased spatial resolution achieved in this study significantly improved our ability to study coherence of gamma oscillations not only between the CA1 and CA3 subfields, but also within these regions. This critical capacity allowed to detect differences in how signals propagate in the hippocampus of wildtype and *Ank3* mutant mice that may have important consequence in the study of bipolar disorder.

Secondly, we expanded the age of the animals used in this study. Many previous studies of gamma band oscillations utilize hippocampal slices from the brains of young animals (REF) but also Hajos et al.((Hájos et al., 2009)) that used animals of just slightly younger in age to our study, while other groups like Gloveli et al. utilized older mice by utilizing a transient induction of gamma oscillations(Gloveli et al., 2005). The use of neonatal animals increases the viability of *ex vivo* slices and can increase the likelihood of acquiring high quality recordings. Generating gamma oscillations in *ex vivo* hippocampal slices requires very healthy FSIs, thus maintaining tissue viability is particularly important. We incorporated the NMDG recovery method in to our approach to enhance the viability of our brain sections (Ting et al., 2014).

The use pMEAs can also help understand how oscillatory activity is generated and sustained at an age where parvalbumin interneurons are more mature. This will prove crucial for further understanding how these cells contribute to the behaviors observed in mouse models of diseases that manifest later in life such as bipolar disorder or schizophrenia. In Nelson et al 2018, we demonstrated that parvalbumin interneurons were greatly impacted by a mutation in the *Ank3* gene. Our preliminary report identified a decrease in inhibitory neurotransmission that altered hippocampal gamma oscillations. Interestingly here we report that the mutant mice exhibit stronger coherence in their gamma oscillations than their wild type littermates.

Previous studies utilizing interface chamber have relied on a combination of kainate and cholinergic agonists to generate the oscillations in various cortical regions such as somatosensory, auditory, and prefrontal cortex, many of which could be relevant to different aspects of the pathology of different diseases. Our novel assay utilizing pMEAS can be adapted to study these various regions with only minor modifications to our current set up. Focusing on these regions could be used to provide additional insight into how the power in gamma propagates through the different cortical layer or to adjacent columns in the case of brain regions like the barrel cortex.

Although we utilized kainate to chemically induce the oscillations, the perforated MEA could be utilized in combination with other methods to induce oscillations such as the local kainate application(Gloveli et al., 2005; McNally et al., 2011) or through selective activation of parvalbumin expressing FSIs using optogenetics (Butler et al., 2018; Cardin et al., 2009; Siegle et al., 2014). Combining our approach with these focalized techniques for generating oscillations could be used to provide a detailed understanding of how oscillatory activity propagates through hippocampus.

One limitation of our study is that we only used adolescent mice aged PND 21-40. Although we did not examine tissue from older animals, we expect our approach could be used to examine gamma oscillations in tissue from older aged animals with minimal adaptation. Modifying these steps could further extend the tissue such as those that would be relevant to diseases with late onset, such as Alzheimer’s disease. Another concern of our approach is the high perfusion rate that we used to provide sufficient, recycling of the bath solution might reduce the amount of waste generated or decrease consumption of precious or expensive compounds.

In summary, we have presented a novel assay that employs perforated MEAs to generate and sustain gamma oscillations in juvenile mice. The high-density geometry of MEA chips allowed us to simultaneously examine the coherence within and between CA1 and CA3 regions. This assay was successfully employed with a mouse model of bipolar disorder revealing an overall increase in coherence despite a decrease in the power in the gamma band. The relevance of this may be pursued further in future studies to increase our understanding of how FSIs dysfunction might be contributing to the pathology of mental health disorders like bipolar disorder.

