## Supplemental Figure 1 for "A Novel Approach to Study Gamma-Band Coherence in *Ex Vivo* Hippocampal Slices from a Mouse Model of Bipolar Disorder"

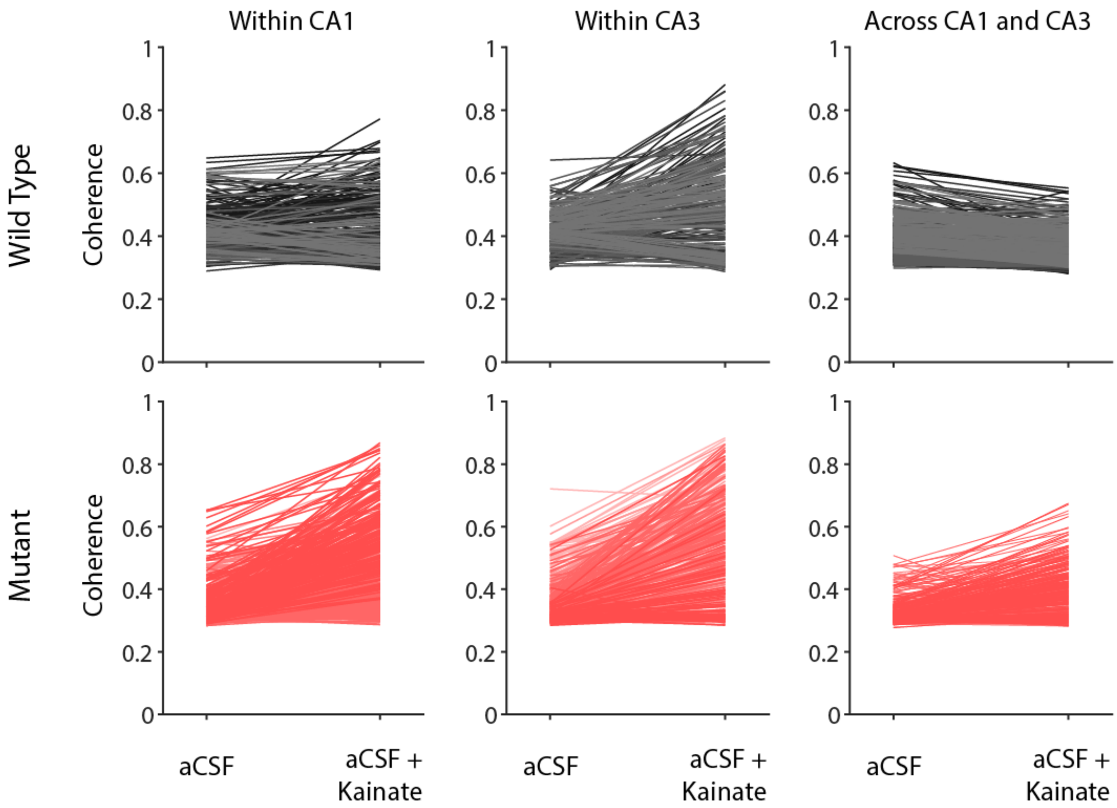


**Figure S1: Coherence increases in individual channels after kainate treatment in the wild type and mutant mouse model of bipolar disorder.** Coherence in individual electrode during the last 5 minutes of aCSF and the last five minutes of aCSF + Kainate (400 nM) for wildtype (top row) and mutant mice (bottom row). Individual electrodes are represented as a line. Gray tones represent electrodes from sections obtained from wildtype mice, while red tones represent those from mutant mice. Each tone represents electrodes from the same slice.
